# Tissue-resident memory T cells mediate mucosal immunity to recurrent urinary tract infection

**DOI:** 10.1101/2021.10.28.466224

**Authors:** Matthieu Rousseau, Livia Lacerda Mariano, Tracy Canton, Molly A Ingersoll

**Affiliations:** Mucosal Inflammation and Immunity, Department of Immunology, Institut Pasteur, 75015 Paris, France and Inserm U1223, Paris France; Université Paris Cité, Institut Cochin, INSERM U1016, CNRS UMR 8104, Paris, 75014, France

## Abstract

Urinary tract infection (UTI) is one of the most prevalent human bacterial infections. New therapeutic approaches, including vaccination and immunotherapy, are urgently needed to combat the rapid global dissemination of multidrug-resistant uropathogens. Development of therapies is impeded by an incomplete understanding of memory development during UTI. Here, we found that reducing bacterial load early in infection, by reducing the inoculum or with antibiotics after infection, completely abrogated the protective memory response. We observed a mixed T helper cell polarization, comprised of Th1, Th2, and Th17 T cells, among T cells infiltrating the bladder during primary infection. Thus, we hypothesized that reducing antigen load altered T helper cell polarization leading to poor memory. Surprisingly, however, T helper cell polarization was unchanged in these scenarios. Instead, we uncovered a population of tissue resident memory (T_RM_) T cells that was significantly reduced in the absence of sufficient antigen. Demonstrating that T_RM_ cells are necessary for immune memory, transfer of lymph node or spleen-derived infection-experienced T cells to naïve animals did not confer protection against infection. Supporting that T_RM_ cells are sufficient to protect against recurrent UTI, animals depleted of systemic T cells, or treated with FTY720 to block memory lymphocyte migration from lymph nodes to infected tissue, were equally protected compared to unmanipulated mice against a second UTI. Thus, we uncovered an unappreciated key role for T_RM_ cells in the memory response to bacterial infection in this mucosa, providing a target for non-antibiotic-based immunotherapy and/or new vaccine strategies to prevent recurrent UTI.

**One Sentence Summary:** T_RM_ are necessary and sufficient for memory to recurrent UTI

## Introduction

Development of immune memory, naturally after infection or by immunomodulatory approaches such as vaccination, protect an organism from experiencing the same infection more than once. Failure to develop adequate immune memory results in repeated or recurrent infection. Mucosal surfaces, which are contiguous with the environment, are particularly at risk for infection.

The bladder is one such mucosal surface that, in a substantial proportion of the population, will be infected repeatedly, often with the same uropathogen (*1*). Indeed, urinary tract infection (UTI), caused predominantly by uropathogenic *E. coli* (UPEC), is one of the most prevalent human bacterial infections (*2*). Women experience UTI at a significantly higher rate than men; and children, the elderly, pregnant women, and immuno-compromised individuals are at greater risk than the general population for complications such as chronic infection, pyelonephritis, and kidney damage (*3–8*). Approximately one in two women will have a UTI in her lifetime, with the highest risk between the ages of 16 and 35, when women are 35 times more likely than men to have a UTI (*9*). Nearly half of all infected individuals will have a recurrent infection, defined by similar or identical symptoms, within 6 months of their first UTI (*10*), suggesting that immune memory fails to develop following infection of the bladder. Although the incidence of UTI is lower in adult men (∼1%), in this population, it is always considered a complicated infection, with considerable risk of becoming a recurrent or chronic UTI (*11*). Finally, the prevalence of UTI increases substantially in older individuals, bringing with it increased morbidity and mortality in an aging population (*12*).

Given the breadth and diversity of at-risk populations, the frequency of recurrence, and the considerable increase in multi-drug resistant uropathogen isolates in global circulation, new approaches, such as immune modulation to augment mucosal immunity, are needed to defend against chronic or recurrent UTI (*13, 14*). First line antibiotic therapy only reduces acute bacterial burden, and no treatment or therapy definitively protects against recurrent UTI (*14*). Indeed, experimental vaccination in humans and mice was reported to induce antibody production correlating with lower bacterial colonization more than 30 years ago, however, these attempts have not lead to a successful clinically available vaccine (*13, 15-21*). Recurrent UTI likely arises due to several factors, including inadequate development of memory responses after a primary infection, but also due to development of reservoirs in the bladder and gut, which potentially escape elimination by antibiotics (*22–24*). As the associated costs of UTI, including treatment and lost productivity, are more than $3 billion in the US (*25*), with similar costs in Europe, a better understanding of how immune memory arises and can be targeted to improve the adaptive response to UTI would have a profoundly positive societal and economic impact, and improve the quality of life of those suffering from recurrent infection.

Notably, how adaptive immunity develops in response to bladder infection is poorly understood (*14, 26*). Urine, from women with a history of UTI, inhibits UPEC binding to human urothelial cells and antibody depletion of this urine abrogates this inhibitory capacity (*27*), suggesting antibodies may protect against recurrent infection. An early key observation, that mice infected with an ovalbumin-expressing UPEC strain have ovalbumin-specific antibodies in the serum and reduced bacterial burden following a second or challenge infection, demonstrates that specific humoral memory can develop during UTI (*28*). We reported that bladder bacterial burden following a challenge infection is lower than that observed during a first or primary UTI (*22*). Bacterial burden is not reduced after challenge infection in dendritic cell-depleted mice, RAG2^-/-^ mice, or in mice depleted of CD4^+^ and CD8^+^ T cells before a primary infection (*22*). Together, these data support that non-sterilizing memory to UTI, mediated by the adaptive immune system, does develop, however, whether T or B cells are the key players in bacterial clearance in the bladder is unclear.

A recent study suggested that a Th2-biased T cell response develops during UTI to promote bladder urothelial repair at the expense of bacterial clearance (*29*). Importantly, while this is seemingly a potential explanation for why the memory response to UTI is nonsterilizing, this work fails to consider that this bias is not observed in male mice even though female and male mice exhibit the same degree of nonsterilizing protection against a challenge UTI (*30*). Indeed, the early cytokine response to UTI, which is robust and diverse in female mice and largely absent in male mice, suggests that a Th2 T cell bias is not the determinant factor in memory development after UTI (*22, 30*).

Thus, we investigated the memory response that arises following UTI, finding that a mixed Th1, Th2, and Th17 bias developed during the first and second infection. Antigen persistence was necessary for immunity, as memory was abrogated after early antibiotic treatment or when mice were infected with a smaller inoculum. Surprisingly, limiting antigen persistence did not change T cell polarization, but did lead to a significant reduction in a population of T cells, identified as tissue resident memory T (T_RM_) cells. Depleting T cells from circulation or blocking lymphocyte egress from lymph nodes before a second infection did not impair immune memory, supporting that local mucosal memory responses, and not systemic immunity, determine protection against UTI, revealing a specific target for new therapeutics.

## Results

### CD4^+^ or CD8^+^ T cells are sufficient for immune memory to UTI

C57BL/6 mice have significantly lower bacterial burdens in their bladders after a second or challenge infection compared to a primary infection with UPEC (*22, 30, 31*). Compared to this control scenario, bacterial burdens are not significantly reduced after a challenge infection in the absence of B or T cells, such as in RAG2^-/-^ mice or in mice depleted of both CD4^+^ and CD8^+^ T cells before the first infection (*22*). Thus, whether B cells or T cells are the key players in bacterial clearance in the bladder is unclear.

To determine the key memory population(s), first, we intravesically instilled 6 to 8-week-old wildtype female C57BL/6 mice or µMT^-/-^ mice, which lack mature B cells (*32*), with 10^7^ colony forming units (CFU) of one of two isogenic UPEC UTI89 strains (UTI89-GFP-amp^R^ or UTI89-RFP-kana^R^). Half of the mice from each group were sacrificed 24 hours post-infection (PI) to assess the bacterial burden in the bladder after the primary infection. Resolution of infection was monitored in the remaining mice by urine sampling twice per week for 4 weeks. The sustained absence of bacteria in the urine was considered to be a resolved infection. At 4 weeks PI, to model a recurrent UTI, all resolved mice were challenged with 10^7^ CFU of the isogenic UPEC strain not used for the primary infection. We sacrificed mice 24 hours post-challenge infection to assess bacterial CFU in the bladder. As expected, in wildtype mice, bacterial burden was significantly reduced 24 hours post challenge (2°) compared to the burden following primary infection (1°). Surprisingly, a comparable, statistically significant reduction in CFU was also observed in µMT^-/-^ mice after challenge infection, supporting that B cells are dispensable for immune memory in UTI (**Fig. 1A**).

**Figure 1.**
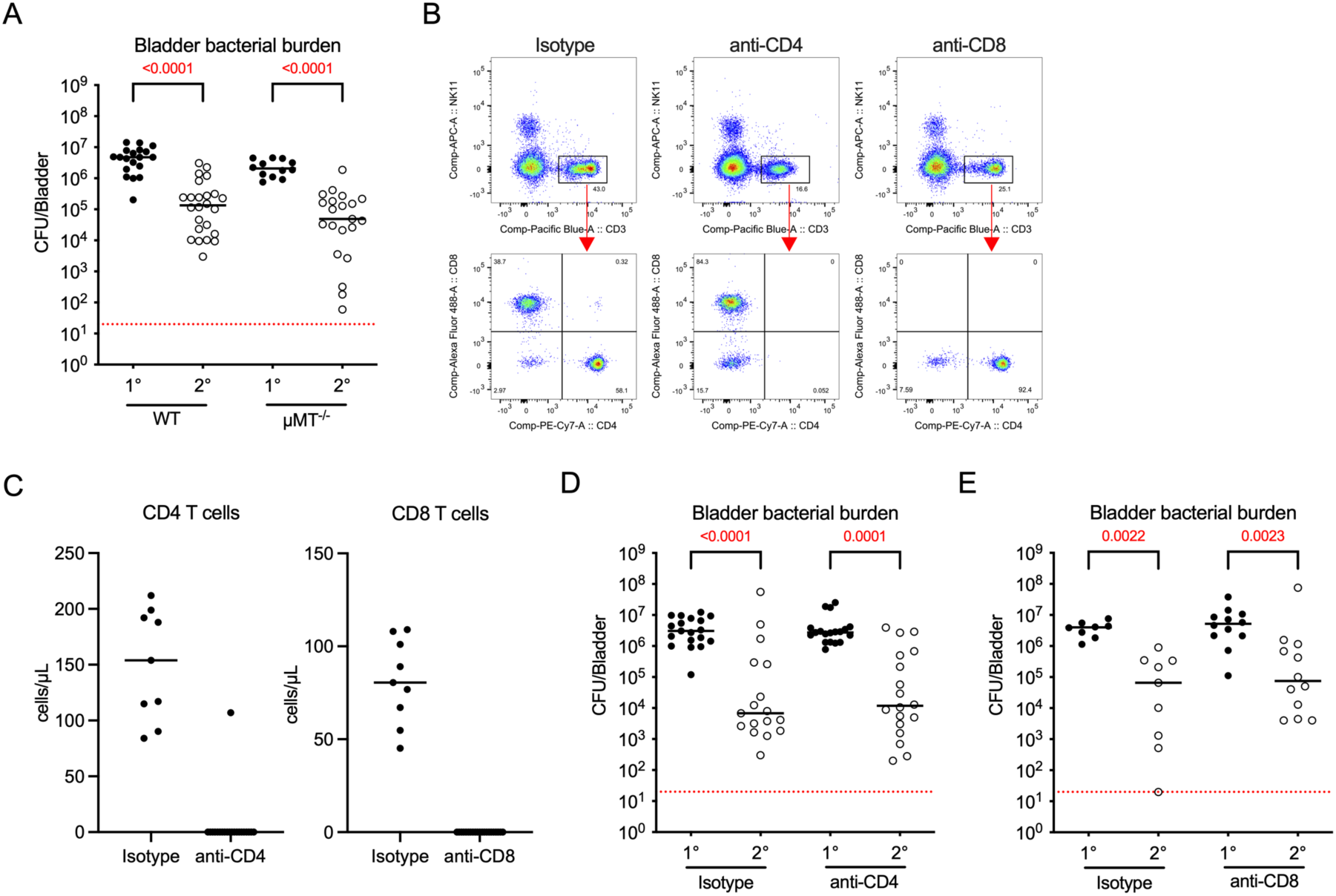
CD4+ or CD8+ T cells can mediate protection against a second UTI. Six- to eight-week-old female C57BL/6 or µMT-/- mice were instilled intravesically with 107 CFU of UPEC strain UTI89 and bladder bacterial burden was assessed 24 hours post-primary infection (1°, filled circles). Urine was monitored twice per week for 28 days and all resolved mice with sterile urine were challenged with an isogenic UTI89 strain and bacterial burden was assessed 24 hours after challenge infection (2°, open circles). (**A**) Graph depicts 1° and 2° CFU per bladder in C57BL/6 (WT) or µMT-/- mice. (**B-E**) Six- to eight-week-old female C57BL/6 mice were treated with anti- CD4 or anti-CD8 depleting antibodies or isotype controls three days prior to infection and again 4 days post-primary infection. Urine was monitored twice per week and all resolved mice were challenged with an isogenic UTI89 strain 28 days later. (**B**) Representative dot plots of circulating blood leukocytes, gated on all CD45+ cells (top row) and CD45+CD3+ cells (bottom row), showing isotype, anti-CD4, or anti-CD8 antibody-treated mice. (**C**) Graphs show quantification of CD4+ and CD8+ T cell depletion in blood. (**D, E**) Graphs depict 1° and 2° CFU per bladder in (**D**) isotype or anti-CD4 antibody-treated mice, or (**E**) isotype or anti-CD8 antibody-treated mice. Data are pooled from 2-5 experiments, n=2 to 7 mice/group in each experiment. Each circle is a mouse, lines are medians. Dotted red lines depict the limit of detection of the assay, 20 CFU/bladder. Significance (adjusted *p*-value) was determined using a Kruskal-Wallis test comparing bacterial burden 24 hours post 1° to 2° within a treatment group, with a Dunn’s post hoc test to correct for multiple comparisons. *p*-values <0.05 are in red.

As we reported that depletion of both CD4^+^ and CD8^+^ T cells before primary infection abrogated memory (*22*), we next tested which T cell subset (CD4^+^ or CD8^+^) was necessary for immune memory to UTI. Six- to eight-week-old female C57BL/6 mice were treated with anti-CD4 or anti-CD8 depleting antibodies or relevant isotype controls (**Fig. 1B, C**), and then infected three days later with 10^7^ CFU of UPEC. As above, bacterial burden was assessed in half of the animals at 24 hours PI and the remaining mice were challenged with 10^7^ CFU of the isogenic UPEC strain following resolution of infection at 28 days PI. In the isotype control groups, bacterial burdens 24 hours after challenge infection were statistically significantly reduced compared to primary UTI, as expected. Unexpectedly, bacterial burdens were also significantly reduced in mice treated with anti-CD4 or anti-CD8 depleting antibodies (**Fig. 1D, E**). Together, our findings support that B cells are dispensable for the adaptive immune response to UTI and either CD4^+^ or CD8^+^ T cells are sufficient for immune memory to infection.

### T cell polarization is mixed during primary and challenge UTI

Having determined that either CD4^+^ or CD8^+^ T cells were sufficient for immune protection, we next wanted to examine early T cell infiltration and accumulation during primary and recurrent UTI. We focused on CD4^+^ T cells because CD4^+^ T cell infiltration was more than 100 times higher than CD8^+^ T cell infiltration following infection of naïve mice (**Fig. S1A**). Following infection with 10^7^ CFU of UPEC, six-week-old female C57BL/6 mice were sacrificed at days 1, 3, and 7 PI or allowed to resolve, infected with the isogenic UIT89 strain 28 days later and sacrificed 1, 3, or 7 days post challenge infection. Total CD4^+^ T cells increased approximately 10-fold in the bladder over one week after primary infection (**Fig. 2A**). Interestingly, T cell numbers in the bladder remained significantly elevated over naïve levels up to one month PI in resolved (R) animals, and increased further 7 days after challenge infection (**Fig. 2A**).

**Figure 2.**
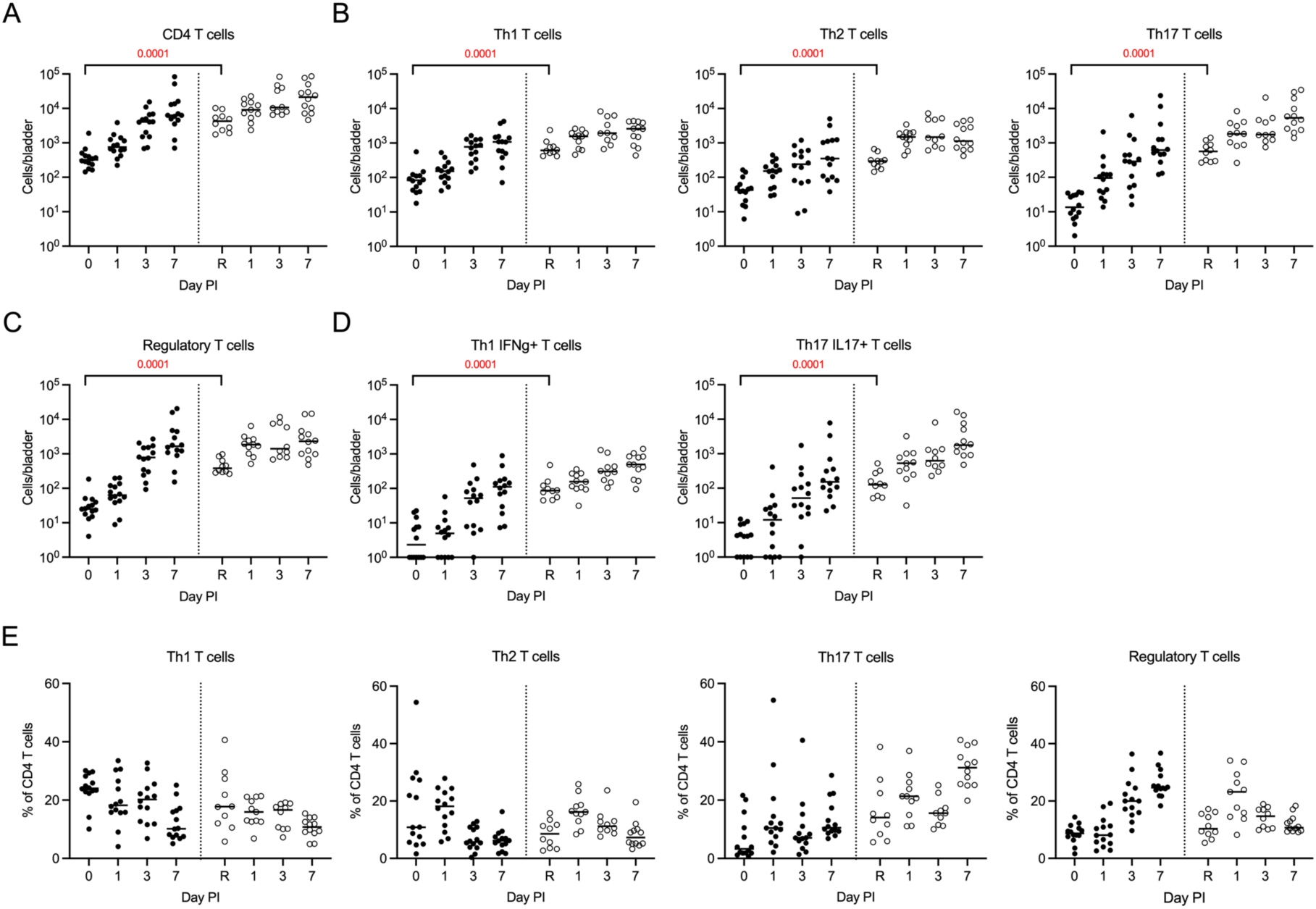
No distinct CD4+ Th cell bias is apparent during primary or recurrent UTI. Six- week-old infected female C57BL/6 mice were instilled intravesically with 107 CFU of UPEC strain UTI89 and sacrificed at the indicated timepoints post-primary (filled circles), or post- challenge infection (open circles). Day 0 are naïve mice and ‘R’ denotes animals that resolved their primary UTI but were not challenged with a second infection. The graphs depict (**A**) total CD4+ T cells, (**B**) total Th1, Th2, and Th17 T cells, (**C**) total Treg T cells, (**D**) total IFNg+ Th1, IL-17+ Th17 T cells, (**E**) percentage of the global CD4+ T cell population for the indicated Th cell subset at the indicated day post primary or challenge infection. The gating strategy is shown in **Fig. S1B**. Data are pooled from 2 experiments, n=5 to 7 mice/group in each experiment. Each circle is a mouse and lines are medians. Nonparametric Mann-Whitney tests comparing cell numbers between ‘0’ and ‘R’ were performed. All *p*-values were corrected for multiple comparisons across all populations shown using the false discovery rate (FDR) method and *p*- values <0.05 are in red.

It was recently proposed that a Th2-polarized immune response mediates nonsterilizing memory to UTI in female mice (*29*). However, female mice express a variety of cytokines in the first 24 hours of UTI that would support type 1, 2, and 3 immune polarization (*22, 30*). Thus, we assessed bladder-infiltrating CD4^+^ T cell polarization at early timepoints after primary and challenge UTI. Using intracellular staining and flow cytometry, we determined T cell polarization by transcription factor and cytokine expression (T-bet^+^, IFNg^+^: Th1; Gata-3^+^, IL-4^+^: Th2; RORgT^+^, IL-17^+^: Th17 (**Fig. S1B, gating strategy**). Reflecting the total numbers, all measured T helper cell subsets increased over time after a first infection and were significantly elevated in infection-experienced, resolved mice (**Fig. 2B**). We also observed that a large number of FoxP3^+^ regulatory T cells (Tregs) infiltrated the bladder after primary UTI (**Fig. 2C**). IFNg^+^ Th1 and IL-17^+^ Th17 subsets were present, but we detected no IL-4- expressing Th2-biased T cells (**Fig. 2D**), in line with our previous finding that IL-4 levels are 10-100 times lower than any other cytokine measured in the bladder (*30*). Of the global CD4^+^ T cell population, Th1 T cells made up ∼10-20% of cells, Th2-polarized T cells represented ∼5-15% of cells, Th17 T cells made up ∼3-30% of cells, and Tregs made up 10-30% of cells over time in primary and challenge infections, and no T helper cell subset dominated the response proportionally (**Fig. 2E**). We observed a similar mixed bias among T cells in the draining lymph nodes after primary and challenge UTI with increased numbers of Th2, Th17, IFNg^+^ Th1, and IL-17^+^ Th17 T cells in resolved mice compared to naïve animals (**Fig. S2A-C**). Together, these data support that, in contrast to a previous report, a dominant T helper bias does not develop during UTI. Interestingly, all CD4^+^ Th cell subsets that increased in the bladder during the first UTI remained significantly elevated over naïve levels in resolved mice, suggesting that infiltrating T cells may undergo tissue-specific imprinting after a primary UTI.

### Antigen persistence is necessary for development of adaptive immunity during UTI

We hypothesized that the abundant Treg infiltration may be indicative of a suppressive or tolerant microenvironment in the bladder in response to UTI. As tolerance can be induced following suboptimal (too much or too little) antigen stimulation (*33, 34*), we tested whether changing the bacterial inoculum would impact development of memory. We previously reported that increasing the inoculum from 10^7^ CFU/mouse to 10^8^ or 10^9^ CFU/mouse does not promote a stronger memory response to a second UTI, supporting that 10^7^ CFU/mouse is sufficient antigen to induce an adaptive immune response (*22*). It may, however, be too high, leading to some degree of tolerance. Indeed, high dose H56 tuberculosis vaccine is detrimental for specific CD4^+^ T cell homing to the lung and protection against tuberculosis, and in some infection scenarios, reducing antigen load leads to better memory development (*35, 36*). Thus, we tested whether decreasing the inoculum, to reduce antigen load, would induce a stronger memory response. We intravesically infected female C57BL/6 mice with 10^7^, 10^5^, or 10^3^ CFU of UPEC. One cohort of mice from each group was sacrificed 24 hours PI to assess bladder bacterial burden and a second cohort was monitored for resolution over 4 weeks. Resolved animals were challenged 28 days later with 10^7^ CFU of the isogenic UPEC strain and we assessed CFU in the bladder 24 hours post-challenge infection. Mice infected with 10^5^ or 10^3^ CFU resolved their primary infection faster than those infected with 10^7^ CFU (**Fig. 3A**). Notably, compared to the control infection with an inoculum of 10^7^ CFU, mice infected with 10^5^ CFU had a productive primary infection but were no longer protected against a challenge infection (**Fig. 3B**). Mice infected with 10^3^ CFU during their primary infection had markedly reduced primary bacterial burdens near or at the limit of detection of the assay. After a challenge infection, these mice were not protected and had bacterial burdens closely resembling the level observed after infection of naïve mice (**Fig. 3B**). Together, this supports that immune memory requires robust bacterial colonization in the bladder and low-level exposure to UPEC cannot induce immune memory.

**Figure 3.**
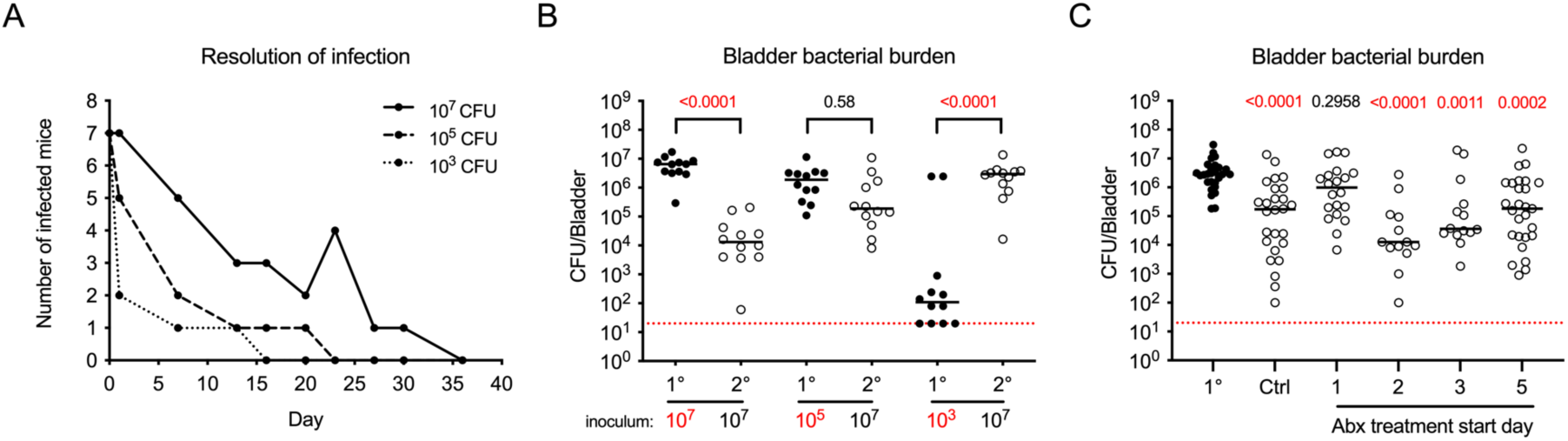
Antigen persistence is necessary for protection against recurrent UTI. 6 to 8-week- old female C57BL/6 mice were infected with (**A, B**) 107 (solid line), 105 (dashed line), or 103 CFU (dotted line) of UPEC strain UTI89 and challenged after resolution with 107 CFU of an isogenic strain 28 days PI, or (**C**) infected with 107 CFU UPEC strain UTI89, treated with trimethoprim/sulfamethoxazole (in drinking water 5 days) starting on days 1, 2, 3, or 5 PI, and challenged with 107 CFU of an isogenic UTI89 strain after resolution on day 28 PI. (**A**) Resolution after primary infection was monitored twice per week by measuring the presence of UPEC in the urine. The graph depicts the number of infected mice over time. An increase in the number of infected mice in the course of the experiment is due to a spontaneous recurrent UTI. (**B, C**) Mice were sacrificed 24 hours post-primary (1°, filled circles) or post-challenge infection (2°, open circles) for CFU assessment. The graphs show bladder bacterial burdens following (**B**) primary infection with the indicated inocula and challenge infection with 107 CFU of UPEC or (**C**) primary and challenge infection in mice treated with trimethoprim/sulfamethoxazole at the indicated days post-primary UTI. The control (Ctrl) group did not receive antibiotics. **A** depicts representative data from 1 experiment of 2 (n=7 mice/group). Data in **B** are pooled from 2 experiments, n=5 to 7 mice/group in each experiment. Data in **C** are pooled from 2-4 experiments, n=6 to 7 mice/group in each experiment. Each circle is a mouse, lines are medians, dotted red lines depict the limit of detection of the assay, 20 CFU/bladder. In **B,** significance was determined by Kruskal-Wallis test comparing bacterial burden 24 hours post 1° to 2° within each inoculum group and in **C**, each 2° group was compared to the 1° CFU, with Dunn’s post hoc test to correct for multiple comparisons and adjusted *p*-values for each comparison are presented, *p*-values <0.05 are in red.

To directly test whether antigen persistence was necessary for immune memory to UTI, we infected mice with 10^7^ CFU of UPEC and treated cohorts of mice with antibiotics at specific days PI. In this scenario, all animals initially have the same antigen exposure to 10^7^ CFU necessary to induce an immune response, but antigen persistence is disrupted at different timepoints PI. The advantage of this approach is that antibiotics are first line therapy to treat UTI, and thus, we modeled the impact of a clinical scenario on immune memory. UPEC-infected female C57BL/6 mice were given trimethoprim/sulfamethoxazole in the drinking water for 5 days starting at 1, 2, 3, or 5 days PI. This antibiotic combination is commonly prescribed for UTI, excreted in the urine, and reported to not change microbiome diversity in humans taking it prophylactically for UTI (*37*). All resolved animals were challenged with an isogenic UPEC strain 28 days PI and bacterial burdens at 24 hours post-challenge infection from all groups were compared to CFU at 24 hours post-primary infection. When antibiotics were given at day 1 post-primary infection, immune memory was abrogated 28 days later, similar to that observed when an inoculum of less than 10^7^ CFU was used for infection (**Fig. 3C**). When antibiotics were delayed until day 2, 3, or 5 post-primary infection, mice developed memory and were protected against challenge infection, similar to that observed in the untreated control group (**Fig. 3C**). These findings support that antigen persistence is necessary for the development of memory in UTI. They also suggest that the loss of protection when antibiotics are given at day 1 PI is not due to nonspecific effects of disruption in the host microbiome, since memory develops when antibiotics are given on day 2 PI. It is important to note that the antibiotic treatment was 5 days in drinking water. Therefore, day 1 treated mice and day 2 treated mice overlapped in antibiotic treatment days by 80% - yet the development of memory was only impacted when antibiotics were given on day 1 PI. Finally, these results raise the important question as to whether a previous UTI is actually a risk factor for recurrent infection (*38*), or rather, early antibiotic treatment compounds to an individual’s risk for recurrent UTI due to poor immune memory.

### Limiting antigen persistence does not impact Th cell polarization

Changing antigen load or persistence abrogated protection against a second infection. Thus, we reasoned that by comparing the T cell response in these infection conditions to the control condition, we could determine mechanisms of immune memory. We first considered that the loss of immune memory observed in the two infection scenarios was due to a change in the proportions of polarized Th cell populations because we previously observed that depletion of tissue resident macrophages before challenge infection results in increased Th1 T cell infiltration and improved bacterial clearance in the bladder (*31*). To test this, we either infected mice with 10^7^ or 10^3^ CFU of UTI89, or we infected mice with 10^7^ CFU of UTI89 and treated half of these mice with antibiotics at day 1 PI. We followed resolution of infection, and 28 days later, we challenged all resolved mice with 10^7^ CFU of UTI89 and assessed CD4^+^ T cell polarization 24 hours post-challenge. In mice infected with 10^3^ CFU for their first UTI, while we observed a significant reduction in the total number of Th1, Th2, and Th17 T cells; IFNg^+^ Th1 T cells, IL-17^+^ Th17 T cells; and Tregs compared to the control group (**Fig. 4A**), all Th cell subsets and Tregs decreased similarly (**Fig. 4B**). Thus, the proportion of each subset and therefore, global Th cell polarization was the same between mice infected with 10^7^ CFU or 10^3^ CFU. In antibiotic-treated mice, we observed no differences in the numbers of Th1, Th2, or Th17 T cell subsets; IFNg^+^ Th1 T cells or IL-17^+^ Th17 T cells; or Tregs compared to untreated control mice (**Fig. 4C**). As the number of T cells in each subset did not change in antibiotic-treated mice compared to the control group, the proportions of each Th cell subset were also not different between the groups (**Fig. 4D)**. Supporting that memory was not lost due to changes in the systemic T cell compartment, the draining lymph nodes also had no differences among any of the parameters measured in the experimental groups compared to the control conditions (**Fig. S3A, B**). Notably, although both experimental groups had abrogated immune memory (**Fig. 3**), in one scenario, T cell infiltration was globally reduced (**Fig. 4A**), whereas in the second, T cell infiltration was not different from the control group (**Fig. 4C**). From these data, we concluded that the loss of immune memory was not due to changes in T cell polarization or potentially even in T cell infiltration after a challenge infection, supporting that the polarization and quantity of infiltrating Th cells in response to UTI are not major determinants of memory development in UTI.

**Figure 4.**
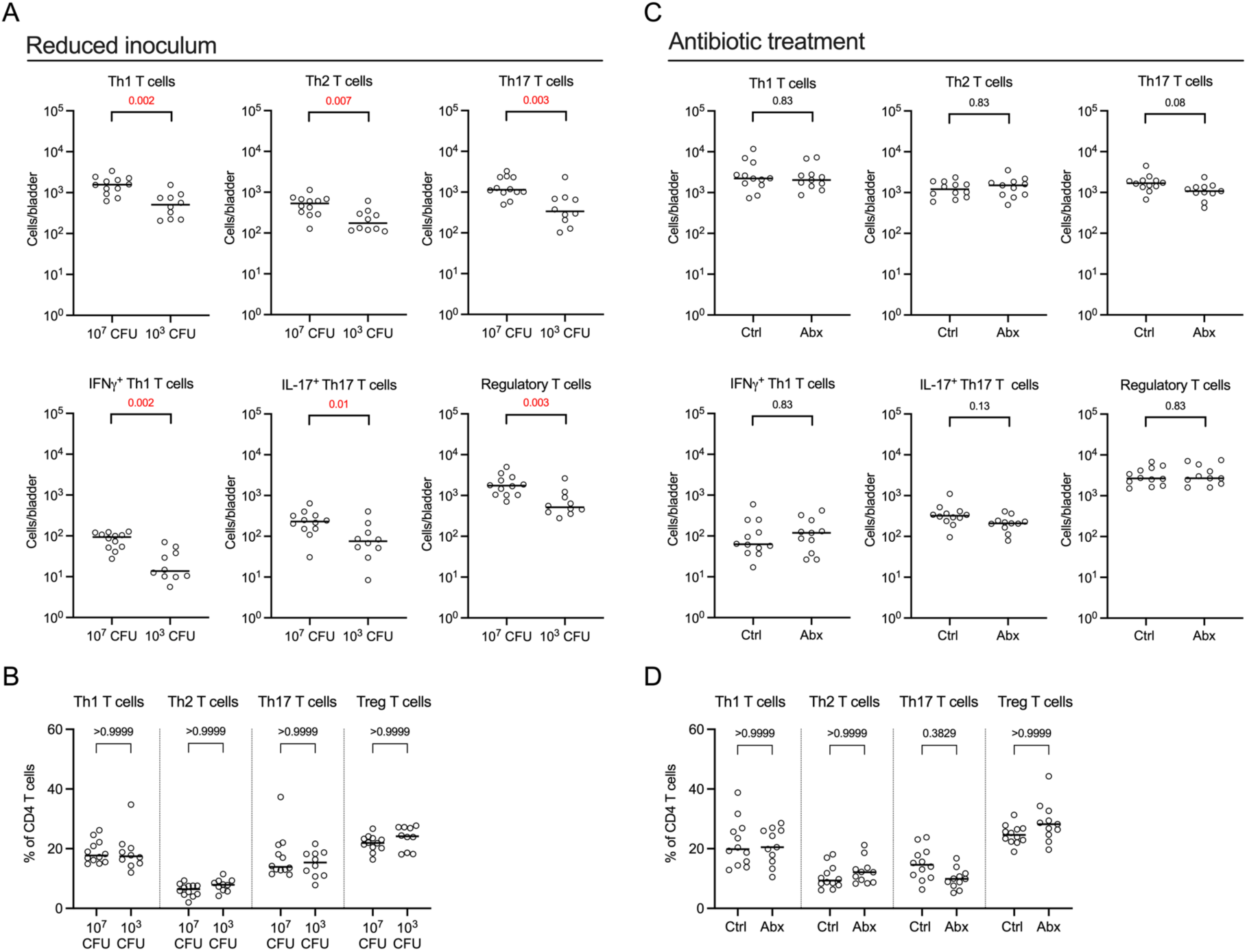
Th T cell polarization is unchanged when antigen persistence is limited. (**A, B**) Six-week-old female C57BL/6 mice were intravesically infected with either 107 or 103 CFU of UTI89, and at day 28 PI, resolved mice were challenged with 107 CFU of UTI89 and bladders analyzed by flow cytometry 24 hours post-challenge infection. Graphs depict (**A**) the total number of the specified Th cell populations per bladder and (**B**) the frequency among CD4+ T cells of the Th cell subsets in **A**. (**C, D**) Six-week-old female C57BL/6 mice were infected with 107 CFU of UTI89. One cohort of mice was treated 24 hours PI with antibiotics and at day 28 PI all resolved mice were challenged with 107 CFU of UTI89. Bladders were analyzed by flow cytometry 24 hours post-challenge infection. Graphs show (**C**) the total number of the specified Th cell populations per bladder and (**D**) the frequency of each Th cell subset among CD4+ T cells. Gating strategy is shown in **Fig. S1B**. Data are pooled from 2 experiments, n=4 to 6 mice/group in each experiment. Each circle represents a mouse and lines are medians. In **A** and **C,** nonparametric Mann-Whitney tests comparing cell numbers between each condition were performed and all *p*- values were corrected for multiple comparisons among the same experiment using the false discovery rate (FDR) method. In **B** and **D,** significance was determined by Kruskal-Wallis tests comparing the frequency of indicated T cell subsets for each condition with Dunn’s post hoc test to correct for multiple comparisons were performed. *p*-values <0.05 are in red.

### Tissue resident memory T cells develop in the bladder after UTI

To understand what cell types are mediating immune memory, we considered that immunological memory can be mediated through B cells, circulating (T_EM_) effector and central (T_CM_) memory T cells, or a more recently described subset of long-lived stationary T cells called tissue resident memory T cells (T_RM_) (*39, 40*). Our prior work, as well as our findings in **Fig. 1**, demonstrated T cells are necessary for an adaptive response while ruling out B cells (*22*). This T cell-mediated memory response to UTI is rapid as a 2-log reduction in bacterial burden is observed in the bladder at 24 hours post challenge infection (*22, 30, 31*). Given the speed of this response, we hypothesized that T_RM_ cells mediate memory to recurrent UTI.

To test this hypothesis, we infected mice with 10^7^ or 10^3^ CFU of UTI89, or we treated mice infected with 10^7^ CFU of UTI89 with antibiotics 1 day PI. 28 days post-primary infection, we challenged all resolved mice with 10^7^ CFU of UTI89 and analyzed bladder resident T cell populations by flow cytometry. Defining these cells as CD3^+^, CD4^+^, CD8^+^, CD44^high^, CD62L^-^, and expressing CD69 and/or CD103, we observed a significant reduction in the number of CD4^+^ and CD8^+^ T_RM_ cells in mice infected with 10^3^ CFU compared to mice infected with 10^7^ CFU (**Fig. 5A**) or in mice treated with antibiotics at 1 day PI compared to control-treated mice (**Fig. 5B**). Supporting that it is specifically bladder-associated T_RM_ cells that mediate memory, there were no differences in the number of T cells with a T_RM_ phenotype in the draining lymph nodes in any of the tested conditions (**Fig. S4A, B**). Notably, in the antibiotic treatment experiment, only T_RM_ cell numbers correlating with a loss of memory, as no other T cell populations were altered in the bladder or draining lymph nodes (**Fig. 4C** and **Fig. S3B**). Thus, reducing antigen load or persistence negatively impacted only local bladder T_RM_ cell populations, correlating with abrogation of protection following a recurrent UTI.

**Figure 5:**
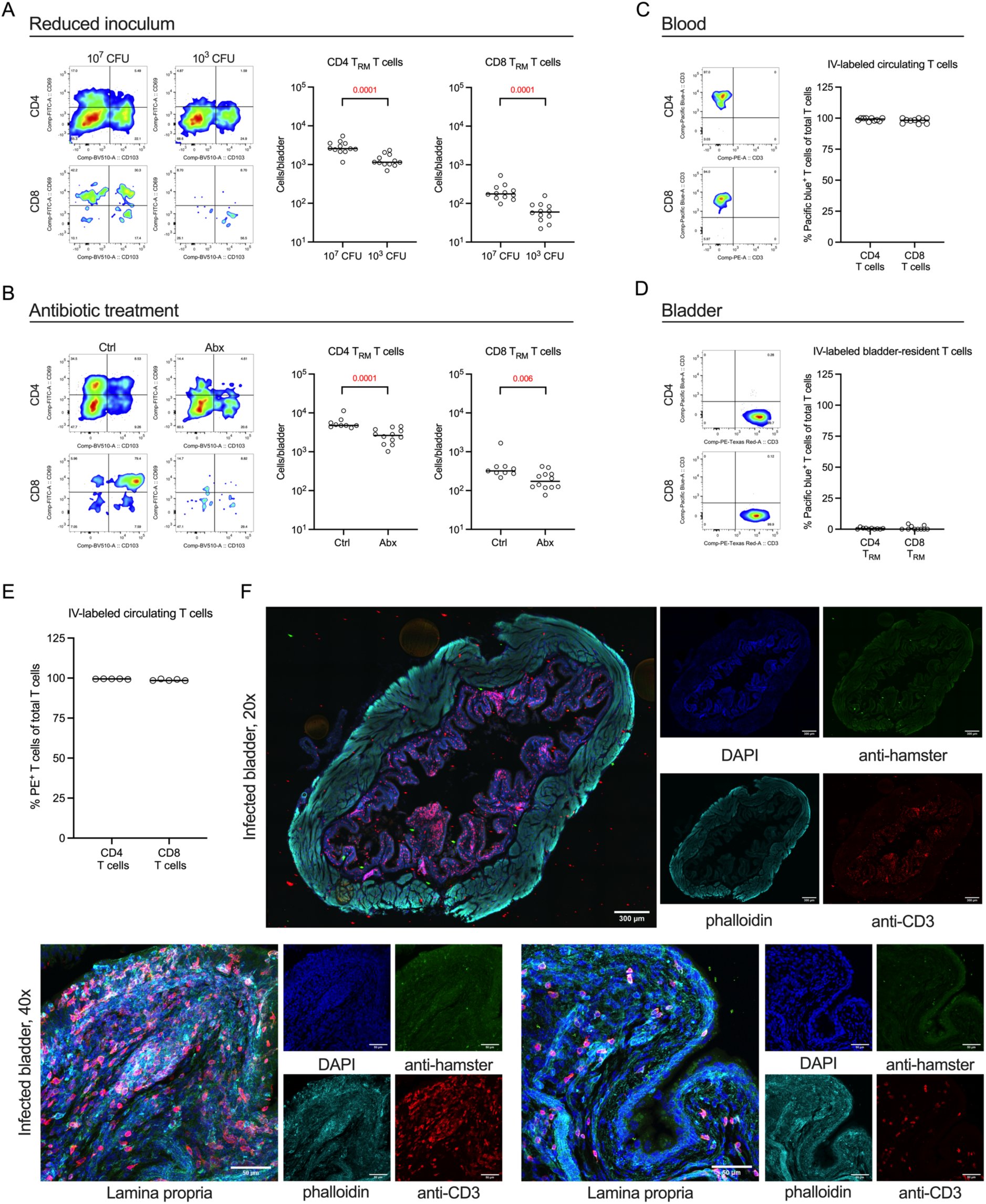
Tissue resident memory T cell accumulation requires antigen persistence. (**A, B**) Representative flow cytometry plots show the phenotype of the specified TRM cell populations quantified in the graphs in bladders after six-week-old female C57BL/6 mice were instilled intravesically with (**A**) 107 or 103 CFU of UTI89 for the primary UTI or (**B**) infected with107 CFU of UTI89 and half of the animals were treated with antibiotics 24 hours post-primary infection. All resolved mice in both scenarios were challenged with 107 CFU of UTI89 28 days later and sacrificed 24 hours post-challenge infection. (**C-F**) Six-week-old female C57BL/6 mice were instilled intravesically with 107 CFU of UTI89 and following resolution 28 days later, received 3µg of anti-CD3 antibody intravenously. Blood and bladders were collected and analyzed by flow cytometry or confocal microscopy. (**C**) Representative dot plots, gated on CD45+CD4+ or CD45+CD8+ cells, and graph show the proportion of IV-injected Pacific blue-labeled anti-CD3 antibody staining on circulating T cells. (**D**) Representative dot plots, gated on CD45+CD4+ or CD45+ CD8+cells, and graph show the proportion of IV-injected Pacific blue-labeled anti-CD3 antibody staining and post-tissue digestion PE-CF594 anti-CD3 antibody staining on bladder- associated T cells. (**E**) Graph shows the proportion of IV-injected PE-labeled hamster anti-CD3 antibody staining on circulating T cells. (**F**) Representative confocal images of 3 different bladders at 20x and 40x magnification. Merged images and single channels with the target of interest are shown. DAPI, 4’,6’-diamidino-2-phenymlindole. Data in **A** and **B** are pooled from 2 experiments, n=3 to 6 mice/group in each experiment. Data in (**C, D**) are pooled from 2 experiments n=5 mice/group in each experiment. (**E**, **F**) are from 1 experiment with 5 mice. Each circle is a mouse, lines are medians. Significance was determined in **A** and **B** by nonparametric Mann-Whitney tests comparing cell numbers between each condition and calculated *p*-values were corrected for multiple comparisons using the false discovery rate (FDR) method. *p*-values <0.05 are in red.

One defining feature of T_RM_ cells is that their tissue resident status renders them inaccessible to intravascular staining strategies. Therefore, to provide supporting evidence that the CD3^+^CD4^+^CD44^high^CD62L^-^, or CD3^+^CD8^+^CD44^high^CD62L^-^ cells expressing CD69 and/or CD103 were indeed tissue resident, we injected fluorescently-labelled anti-CD3 antibody intravenously (IV) after resolution of a primary infection and collected blood and bladders 3 minutes after injection. Circulating CD4^+^ and CD8^+^ T cells were nearly universally labeled by the injected antibody (**Fig. 5C**). By contrast, bladder-associated CD4^+^ and CD8^+^ T cells, including CD4^+^ and CD8^+^ T_RM_ cells, were almost entirely negative for the IV-injected antibody, but were identified by using an anti-CD3 antibody with a different fluorophore after bladder digestion for flow cytometry (**Fig. 5D**). As a complementary approach to test whether these cells were tissue resident while avoiding the caveats of tissue disruption, we used confocal microscopy following IV-labeling of T cells. Again, blood CD4^+^ and CD8^+^ T were uniformly positive for an IV-injected fluorophore labeled anti-CD3 antibody (**Fig. 5E**). However, using an anti-hamster secondary antibody to amplify the IV-injected primary anti-CD3 antibody signal in the bladder, we did not detect any T cells labeled by IV antibody administration (**Fig. 5F**). The absence of signal with anti-hamster labeling was not due to a lack of T cells as when we used an anti-CD3 antibody to immunostain bladder tissue sections, we observed an abundance of T cells in the urothelium and lamina propria of 28 day post-infected tissue (**Fig. 5F**). Altogether, these data support that CD3^+^CD4^+^, CD8^+^, CD44^high^, CD62L^-^ cells expressing CD69 and/or CD103 are T_RM_ cells.

### Tissue resident memory T cells are necessary and sufficient to mediate immune memory to recurrent UTI

To formally demonstrate that T_RM_ cells mediate protection against a recurrent UTI, we hypothesized that their absence should abrogate immune memory and their presence, in the absence of all other memory cells, should confer protection. To address the necessary side of this equation, we would need to eliminate T_RM_ cells. However, there are currently no specific tools that target T_RM_ cells without also impacting circulating T cell populations. To circumvent this challenge, we took an adoptive transfer approach in which we infected mice with UPEC, allowed the animals to resolve, and then transferred CD4^+^ and CD8^+^ T cell populations from the spleen and bladder-draining lymph nodes to naïve mice (**Fig. 6A, B**). To ensure sufficient numbers of T cells, we transferred 10^5^ T cells to a cohort of mice and 10^6^ T cells to a second cohort. We compared the level of protection 24 hours PI in mice receiving central and effector memory T cell transfer to animals that had been previously infected and challenged. In this scenario, only infection experienced mice would have T_RM_ cells in the bladder. We observed that, in contrast to mice that were infected and then challenged, naïve mice receiving 10^5^ or 10^6^ T cells were not protected against UPEC infection, and had similar levels of bacteria in their bladders compared to unmanipulated naïve mice (**Fig. 6C**).

**Figure 6:**
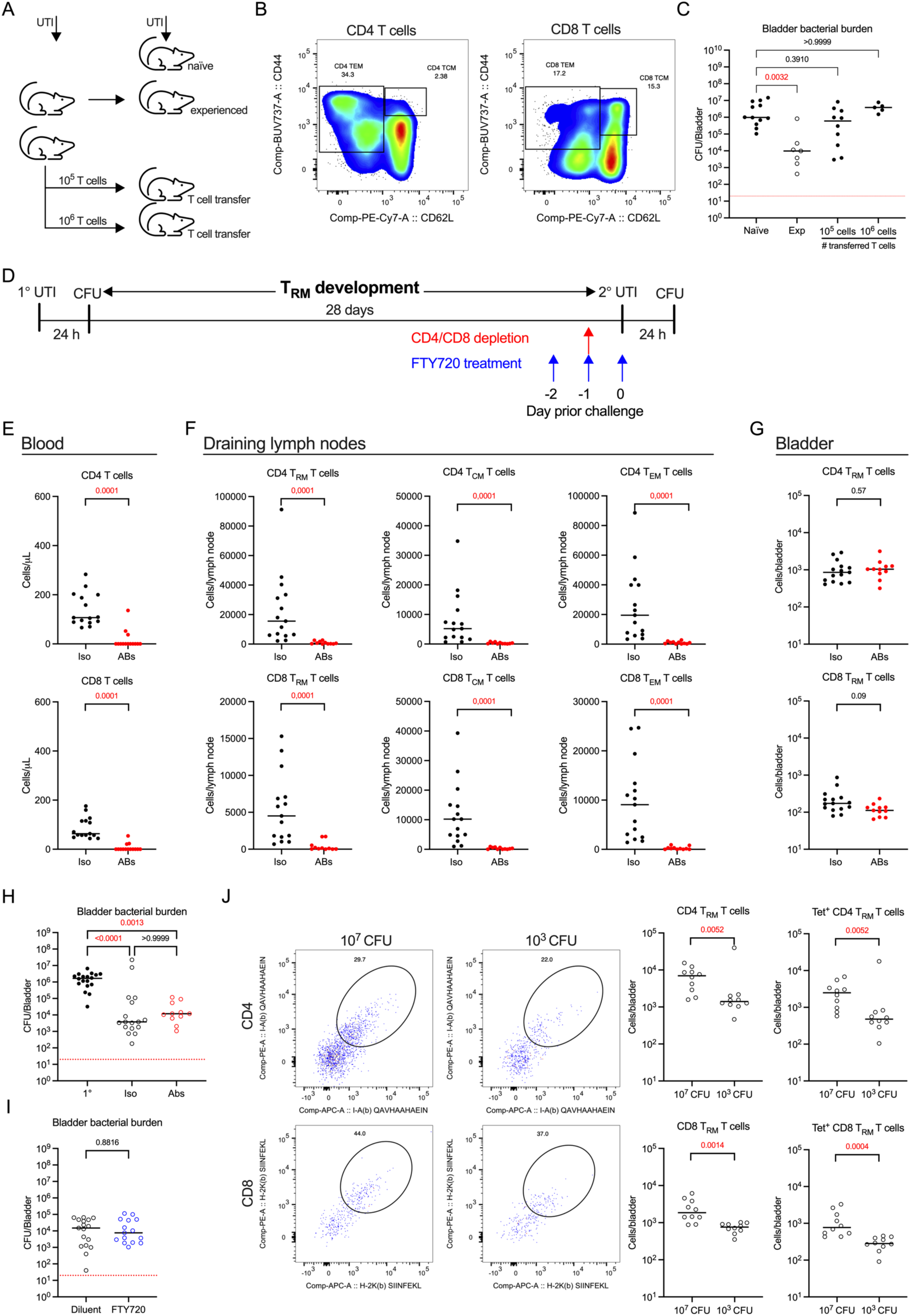
Tissue resident memory T cells mediate protection against recurrent UTI (A) Schematic of experiments in which 10 week old female C57BL/6 mice received T cells by adoptive transfer before infection with 107 CFU of UTI89. (**B**) Representative flow plots show the phenotype of CD4+ and CD8+ T cells transferred from infected and resolved mice to naïve mice prior to infection. (**C**) Graph shows the bacterial burden in naïve mice, infected and challenged mice (exp - experienced), and naïve mice that received 105 or 106 T cells 36 hours prior to UPEC infection. **(D**) Schematic of experiments shown in **E-I**, in which six-week-old female C57BL/6 mice were infected with 107 CFU of UTI89 and at 28 days post-infection treated with either anti-CD4 and anti-CD8 antibodies or their respective isotype controls; or FTY720 or the control diluent. (**E-G**) Graphs depict the total number of specified cell populations in (**E**) blood, (**F**) bladder-draining lymph nodes, and (**G**) bladder in resolved but unchallenged mice sacrificed to assess depletion. (**H, I**) Graphs show bacterial burden in (**H**) control and antibody- depleted mice and (**I**) control and FTY720-treated mice challenged with 107 CFU of UTI89 and analyzed 24 hours post-challenge infection. (**J**) Representative flow cytometry plots depict tetramer staining of CD4+ or CD8+ TRM cells, and graphs show the total number of CD4+ or CD8+ TRM cell populations per bladder, and the number of ova-specific, tetramer+ TRM cells per bladder, after six-week-old female C57BL/6 mice were instilled intravesically with 107 or 103 CFU of UTI89-GFP-ampR-ova, and following resolution, challenged on day 28 PI with 107 CFU of UTI89-GFP-ampR-ova and sacrificed 24 hours post challenge infection. Data in **C, E**-**H** are pooled from 3 experiments, n=3 to 6 mice/group in each experiment. Data in **I** are pooled from 4 experiments, n=3 to 5 mice/group in each experiment. Data in **J** are pooled from 2 experiments, n=5 mice/group. Each circle is a mouse, lines are medians. In **H** and **I**, dotted red lines depict the limit of detection of the assay. Significance in **C** was determined by nonparametric Mann- Whitney tests comparing CFU in the experimental groups to the control condition of naïve mice. Significance in **E-G** was determined by nonparametric Mann-Whitney tests comparing cell numbers between each condition and all calculated *p*-values were corrected for multiple comparisons using the false discovery rate (FDR) method, in **H** by Kruskal-Wallis test comparing bladder bacterial burden 24 hours post primary infection (filled circles) to bacterial CFU 24 hours post challenge infection (open circles) in isotype (black open circles) or depleting antibody- treated mice (red open circles), with Dunn’s post hoc test to correct for multiple comparisons, in **I** by nonparametric Mann-Whitney test comparing bacterial burden 24 hours post challenge between diluent (black open circles) and FTY720-treated mice (blue open circles), and in **J** by nonparametric Mann-Whitney tests comparing cell numbers between each condition and all calculated *p*-values were corrected for multiple comparisons using the false discovery rate (FDR) method. *p*-values <0.05 are in red.

Having demonstrated that T_RM_ cells were necessary for protection against a second infection, we next eliminated all T cells subsets except T_RM_ cells to test whether they were also sufficient for immune memory. Similar to their inaccessibility to intravascular labeling, T_RM_ cells are protected from depleting antibodies when administered using specific doses (*41–44*). Thus, we created a scenario in which the only UPEC-specific memory T cells present during a challenge infection would be T_RM_ cells in the bladder. Mice were infected with UPEC and a cohort of animals was sacrificed 24 hours PI infection. After resolution of infection, at day 28, mice received a single dose of anti-CD4 and CD8-depleting antibodies or their respective isotype controls. Mice were challenged 24 hours later with 10^7^ CFU of UPEC (**Fig. 6D**). In a group of T cell-depleted but uninfected mice, we measured T cell numbers in the blood, draining lymph nodes, and bladder. CD4^+^ and CD8^+^ T cells were significantly reduced to nearly zero in the circulation of antibody-treated mice (depicted in red) compared to control isotype-treated animals (**Fig. 6E**). In bladder-draining lymph nodes of antibody-depleted mice, CD4^+^ and CD8^+^ T cells with a T_RM_ cell phenotype, as well as CD4^+^ and CD8^+^ T_CM_ and T_EM_ cells (**Fig. 6F**) were significantly reduced to nearly zero, indicating depleting antibodies accessed the lymph nodes and any memory T cells resident in the lymph nodes were no longer present to infiltrate the bladder upon challenge infection to mediate protection. Importantly, analysis of the T_RM_ populations in the bladder revealed that T_RM_ cells were protected from antibody depletion and there was no reduction of CD4^+^ or CD8^+^ T_RM_ cells in the bladders of mice that received CD4- and CD8-depleting antibodies compared to mice that were treated with isotype controls (**Fig. 6G**). Thus, only bladder T_RM_ cells were present in the mice when we challenged them with a second UPEC infection.

Following challenge UTI, the control isotype-treated mice had bacterial burdens that were significantly reduced compared to bacterial CFU measured after a primary infection, as expected (**Fig. 6H**). Remarkably, anti-CD4 and CD8 antibody-treated mice were also protected from challenge infection to the same extent as control animals (depicted by red open circles), with significantly reduced bacterial CFU in the bladder despite a near global circulating T cell depletion, demonstrating that immune memory was intact in these animals (**Fig. 6H**). As an alternative approach to reinforce this finding, we used FTY720, an inhibitor of the sphingosine-1-phosphate receptor pathway (*45*), which prevents lymphocyte migration from the lymph nodes. Mice were infected with UPEC, and after resolution of infection at day 28, received 3 doses of either control diluent or FTY720 prior to challenge infection (**Fig. 6D**). Control or FTY720-treated mice (depicted by blue open circles) were protected as well as control-treated mice 24 hours post-challenge with no significant difference in their bacterial burdens after a second UTI (**Fig. 6I**). Together, these results support that T_RM_ cells, specifically in the bladder, and not peripheral memory T cells, mediate protection against recurrent UTI.

Finally, we hypothesized that the T_RM_ cells that control bacteria in a recurrent infection and mediate the memory response were pathogen-specific. To test this hypothesis, we infected mice with either 10^7^ or 10^3^ CFU of a UTI89 strain that also expresses two ovalbumin peptides that are presented in the context of class I and class II and can be followed using MHC I and MHC II tetramers, respectively (UTI89- GFP-amp^R^-ova, OVA257-264 and OVA323-339). 28 days post-primary infection, we challenged all resolved mice with the UTI89 strain expressing the ova peptides and analyzed bladder resident T cell populations by flow cytometry. To detect ova-specific CD4^+^ and CD8^+^ T_RM_, we used I-A(b) QAVHAAHAEIN, and H-2K(b) SIINFEKL tetramers, respectively. Similar to our results in **Fig. 5A**, we observed a significant reduction in the total number of CD4^+^ and CD8^+^ T_RM_ cells in mice infected with 10^3^ CFU compared to mice infected with 10^7^ CFU of UTI89-GFP-amp^R^-ova (**Fig. 6J**). Importantly, ova-specific tetramer^+^ CD4^+^ and CD8^+^ T_RM_ were also significantly reduced in mice infected with 10^3^ CFU compared to mice infected with 10^7^ CFU (**Fig. 6J**). All together, these results support that bacteria-specific T_RM_ T cells accumulate in the bladder after a primary UTI. They require antigen persistence for their development and they are both necessary and sufficient for a memory response to a recurrent UTI.

## Discussion

Overwhelmingly, studies of immunity to UTI focus on innate immunity, and despite how common recurrent UTI is, we have a poor understanding of how memory develops after an acute infection (*14*). Supporting that humoral responses are important, UPEC-specific antibody production is observed in children with pyelonephritis (*46*) and, many years ago, antibodies were proposed as a biomarker for human pyelonephritis (*47*). Thus, it was surprising to rule out a role for B cells in our model of recurrent UTI. Notably, however, in a similar mouse UTI model, antibodies to UPEC arise only when bacteria colonize both the bladder and kidneys (*48*). We used C57BL/6 mice in our studies, and given that this background has a very low incidence of vesicoureteral reflux and kidney colonization (*49*), and that antibodies are present specifically in cases of human pyelonephritis, our findings support that B cells are likely dispensable for memory to cystitis. This has consequences for future vaccines or other immunomodulatory strategies for recurrent UTI. Indeed, it may be that targeting the bladder to promote antibody production will not provide efficacious protection against cystitis, and may explain why, despite more than 25 years of research, vaccination strategies for UTI have so far not been successful.

Immune memory to UTI develops, but it is not sterilizing (*22*). Inadequate or inefficient memory responses likely play a role in the frequency of recurrent UTI. Recently, a study proposed that a type 2 immune biased response during UTI in mice is the reason for why memory is not sterilizing (*29*). While a Th2 biased response would be expected to favor tissue repair over strong cellular immunity, our data show that immune memory can be ablated without changing the proportion, or even the number, of T helper subsets and that there is no prevailing T helper cell bias in the first 7 days of a primary or recurrent infection. In addition, female and male mice develop comparable protective memory responses after an initial UTI, although male mice do not express type 2 cytokines (*30*). In trying to understand this paradox, we found through immunophenotyping that the immune response is not specifically biased towards any single Th subset, but rather that a mixed Th1, Th2, and Th17 response develops, with a robust infiltration of regulatory T cells. While this mixed response was not previously appreciated, it is not entirely surprising, as in the first 24 hours of infection, UPEC induces extensive tissue damage, including robust inflammation and urothelial cell exfoliation, and will be both intracellular and extracellular (*22, 30, 50-53*), necessitating a Th2 response to mediate repair, regulatory T cells to control effector cell activation, Th1-polarized T cells to eliminate intracellular pathogens and a Th17 response to target mucosal and extracellular pathogens concomitantly (*54–58*). Indeed, this mixed response reflects the complexity of the host-pathogen interaction and immune response to UTI, as well as the fine equilibrium that must be established between resolution of infection and maintenance of bladder integrity. How each of these T helper subsets contributes to a future pool of T_RM_ cells after a primary infection will be challenging but interesting to unravel.

Adding to our understanding of memory to UTI, the quantity and persistence of bacteria is a key parameter in the development of an adaptive immune response. A minimum exposure of 10^7^ CFU and of greater than 24 hours is needed for the development of adaptive immunity after a primary UTI, in mice, however, the inoculum size needed to induce memory in humans is unclear. Indeed, whether humans are infected with a “high” or “low” inoculum is a major unknown in the field and challenging to determine. The variable start of antibiotics further complicates understanding how a human infection may progress. One potential caveat of our antibiotic experiment is that trimethoprim-sulfamethoxazole may induce alterations in the mouse microbiome, impacting T_RM_ development. Given that antibiotic treatment only impacted memory when given at 24 hours PI and not when given at 48, 72, or 120 hours later, supports that this is unlikely, however it cannot be definitely ruled out. Indeed, the microbiome can impact T_RM_ development, but the relationship is complex and requires further studies (*59*). While additional studies are necessary to determine the mechanisms underlying these results, alarmingly, it may be that the rapid availability of antibiotics to individuals with a UTI actually places them at risk for recurrent infections by impeding the development of immune memory. Determining whether delaying antibiotic treatment improves the memory response is feasible and may ultimately decrease the incidence of recurrence. Treatments that ameliorate symptoms but do not eliminate bacteria themselves, followed by delayed antibiotic therapy in those not at risk for pyelonephritis, may be a better therapeutic approach for lasting memory. Supporting this idea, while nonsteroidal anti-inflammatory drugs (NSAIDs) administered in place of antibiotics are not superior to antibiotic treatment (*60–62*), a meta-analysis of 962 patients comparing antibiotic treatment to placebo for uncomplicated cystitis did not find a difference in the development of pyelonephritis (*63*). This suggests that delaying antibiotics may not put individuals at greater risk for progression to pyelonephritis, and may support the development of a better memory response. Further supporting this alternative treatment approach, NSAID-treated individuals had a significant reduction in the number of recurrent UTI in their first month post-treatment compared to those treated with antibiotics (*64*). Thus, it would be of great value to test whether briefly delaying antibiotic treatment promotes a better memory response and a reduction in recurrent UTI in humans, which would, in turn, reduce reliance on antibiotics for this infection.

We previously reported that macrophage depletion before a primary or a challenge infection improves bacterial clearance during the challenge infection (*22, 31*). This phenotype is T cell-dependent, and macrophage depletion specifically before a second UTI leads to an increase in type 1-biased effector immune cell infiltration (*22, 31*). Thus, when we observed that memory was negatively impacted by decreasing the inoculum or administering antibiotics, we expected a change in T cell polarization, potentially towards a Th2 bias. We were surprised to observe changes only in the numbers of T_RM_ cell populations, which suggests to us that resident macrophages directly influence bladder T_RM_ cell development or maintenance, potentially as a mechanism to maintain bladder integrity. Bladder resident macrophages likely modulate the microenvironment, which shapes the local T_RM_ cell response. In other mucosal tissues, lung resident macrophages limit the development of influenza-specific CD8^+^ T_RM_ cells (*65*). In *Yersinia pseudotuberculosis* infection, type I IFN and IL-12 in the intestinal microenvironment are key for differentiation and persistence of T_RM_ cells, and depletion of CCR2^+^ IL-12-producing cells, including macrophages or monocytes, impedes T_RM_ cell differentiation (*66*). Deciphering the mechanisms by which macrophages control local T cell responses in the bladder will be key to design immunomodulatory approaches.

The discovery of T_RM_ cells localized in non-lymphoid tissues, which are anatomically well positioned to detect and respond to a second infection faster than T_CM_ or T_EM_ cells, has driven development of vaccine strategies to promote their differentiation. UTI needs new therapeutic approaches to combat the rapid global spread of multidrug resistant bacteria. Vaccination strategies promoting the differentiation of bladder T_RM_ cells would likely be of great benefit for patients suffering from recurrent UTI and would have a positive economic impact by reducing the cost of medical care. The failure of previous vaccine strategies for UTI may be because the correct vaccination strategy to promote the differentiation of T_RM_ cells has not been identified. For example, a BCG mucosal vaccination better protects mice against tuberculosis compared to a subcutaneous vaccination, and adoptive transfer of T_RM_ cells from vaccinated mice protects naïve animals from disease (*67*). Similarly, a mucosal vaccine approach provides superior protection against coronavirus infections compared to a subcutaneous route (*68*). In addition to the route of vaccination, the type of vaccine used is also a key parameter. Indeed, an attenuated influenza vaccine promotes T_RM_ cell generation in the lung, inducing long term protection, whereas an inactivated vaccine does not, despite intranasal delivery (*69*). Intranasal immunization with *Chlamydia trachomatis* promotes uterine mucosa protection and oral or gastrointestinal vaccination with *Chlamydia muridarum* protects the genital tract from infection (*70, 71*). These observations will likely inform vaccine strategies for UTI and suggest that the bladder mucosa does not need to be directly targeted.

In sum, we demonstrated that bladder T_RM_ cells mediate memory to recurrent UTI, and do so in the absence of circulating or lymph node-resident T cells. Thus, we uncovered a specific, targetable population for development of new therapeutic approaches for this common infection plagued by antibiotic resistance. These findings greatly improve our understanding of adaptive immunity to UTI and provide valuable information for the design and development of much needed new non-antibiotic-based therapies.

## Materials and methods

### Study design

This study was conducted using a preclinical mouse model in controlled laboratory experiments to investigate adaptive immunity to UTI. Our objective was to determine how an adaptive immune response develops after a UTI. Mice were assigned to groups by random partition into cages. In all experiments, a minimum of 2 and a maximum of 7 mice made up an experimental group and all experiments were repeated 2 to 5 times, with the exception of confocal imaging, in which five mice were assessed in total. Data from all repetitions were pooled before any statistical analysis to reach a predetermined *n* of at least 6-10 mice per group. As determined *a priori*, all animals with abnormal kidneys (atrophied, enlarged, and/or white in color) at the time of sacrifice were excluded from all analyses, as we have observed that abnormal kidneys correlate with the inability to resolve infection. Animals with a recurrent or unresolved UTI, defined as having persistent bacteria in the urine up to 1 day prior to challenge infection, were also excluded. End points were determined before the start of experiments and researchers were not blinded to experimental groups.

### Ethics Statement

Animal experiments were conducted in accordance with approval of protocol number 2012-0024 and 2016-0010 by the *Comité d’éthique en expérimentation animale Paris Centre et Sud* and the *Comités d’Ethique pour l’Expérimentation Animale* Institut Pasteur (the ethics committee for animal experimentation), and APAFIS #34290 by SC3 - CEEA34 – Université de Paris Cité, at Institut Cochin, in application of the European Directive 2010/63 EU. In all experiments, mice were anesthetized by intraperitoneal injection of 100 mg/kg ketamine and 5 mg/kg xylazine and sacrificed by carbon dioxide inhalation or cervical dislocation after isoflurane inhalation.

### Mice

Female mice between the ages of 6 and 8 weeks were used in this study. Female C57BL/6J mice were obtained from Charles River Laboratories France. µMT ^−/−^ mice were bred in house at Institut Pasteur, Paris, and were a kind gift from Claude Leclerc.

### UTI and determination of bacterial burden

Female mice were anesthetized as described above, catheterized transurethrally, and infected with 10^7^, 10^5^, or 10^3^ colony forming units (CFU) of UTI89-GFP-amp^R^ or UTI89-RFP-kan^R^ in 50 µL PBS as previously described (*22*). UTI89-GFP-amp^R^ and UTI89-RFP-kan^R^ are isogenic UTI89 strains that infect with equal efficiency and were used interchangeably (*22*). For flow cytometry, only the nonfluorescent parental UTI89 strain was used. In the transfer of experienced T cells, and tetramer- staining experiments (**Fig. 6**), mice were infected with 10^7^ or 10^3^ CFU UTI89-RFP-kan^R^-ova or UTI89- GFP-amp^R^-ova. These strains were constructed by integration of the ovalbumin peptides (OVA257-264- GGspacer-OVA323-339) into the bacterial chromosome in our UTI89-GFP-amp^R^ and UTI89-RFP-kan^R^ strains (Recombina Biotech, Navarra Spain). Infection was monitored by detection of bacterial growth from urine samples collected twice per week. 2 µL of urine were diluted directly into 8 µL PBS spotted on agar plates containing antibiotics as appropriate (kanamycin (50 µg/ml) or ampicillin (100 µg/ml)). The presence of any bacterial growth was counted as positive for infection. The limit of detection (LOD) for this assay is 500 bacteria per mL of urine. When indicated, mice were treated with trimethoprim (40mg/kg)/sulfamethoxazole (200mg/kg) in the drinking water for 5 days. Mice were sacrificed at indicated time points, bladders homogenized in sterile PBS, serially diluted, and plated on LB agar plates with antibiotics, as appropriate, to determine CFU. The LOD for CFU in the bladder is 20 CFU per bladder and is indicated by a dotted line in graphs. All sterile organs are reported at the LOD.

### Flow cytometry of bladder tissue, draining lymph nodes and blood

Samples were acquired on a BD LSRFortessa using DIVA software (v8.0.1), and data were analyzed by FlowJo (Treestar) software. Bladder and blood analyses were performed as described previously (*22, 31, 72*). Briefly, bladders were dissected, cut into small pieces, and digested using Liberase (0.34 U/ml) in phosphate-buffered saline (PBS) at 37°C for 1 hour with robust manual agitation every 15 min. Digestion was stopped by adding FACS buffer (PBS supplemented with 2% fetal bovine serum and 0.2 µM EDTA). Single cell suspensions were washed and resuspended in brilliant stain buffer (BD) with anti-mouse CD16/CD32 to block Fc receptors. Antibody mixes (**Table 1**) in brilliant stain buffer were added directly to the samples after 10 minutes. Total cell counts were determined by addition of AccuCheck counting beads to a known volume of sample after staining, just before cytometer acquisition. To determine cell populations in the circulation, whole blood was incubated with BD PharmLyse and stained with antibodies (**Table 1**). Total cell counts per µL of blood were determined by the addition of AccuCheck counting beads to 10 µl of whole blood in 1-step BD Fix/Lyse Solution. Draining lymph nodes were disrupted with 27G needles and passed through 70 µm filters (Miltenyi). Single cell suspensions were washed and resuspended in brilliant stain buffer with anti-mouse CD16/CD32 to block Fc receptors and subsequently stained as above for bladders. Total cell counts were determined as they were for the bladder. Memory T cell staining was performed at 37°C for 30 minutes to enhance CCR7 staining (*73*).

**Table 1:** Antibodies used for flow cytometry in this study.

| Antibodies | Clone | Source | Identifier |
| --- | --- | --- | --- |
| Anti-mouse NK1.1 APC | PK136 | BD Biosciences | cat# 550627 |
| Anti-mouse B220 AF700 | RA3-6B2 | BD Biosciences | cat# 557957 |
| Anti-mouse I-A/I-E (MHCII) APC-eFluor780 | MS/114.15.2 | Invitrogen | cat# 47-5321-82 |
| Anti-mouse CD8 FITC | 53-6.7 | BD Biosciences | cat# 553031 |
| Anti-mouse GR1 PerCP-Cy5.5 | RB6-8C5 | BD Biosciences | cat# 552093 |
| Anti-mouse CD3 Pacific Blue | 500A2 | BD Biosciences | cat# 558214 |
| Anti-mouse CD45 BV605 | 30-F11 | BD Biosciences | cat# 563053 |
| Anti-mouse CD115 PE | AFS98 | Invitrogen | cat# 12-1152-82 |
| Anti-mouse SiglecF PE-CF594 | E50-2440 | BD Biosciences | cat# 562757 |
| Anti-mouse CD4 PE-Cy7 | GK1.5 | ebioscience | cat# 25-0041-82 |
| Anti-mouse NK1.1 AF700 | PK136 | BD Biosciences | cat# 560515 |
| Anti-mouse CD127 APC-eFluor780 | A7R34 | Invitrogen | cat# 47-1271-82 |
| Anti-mouse CD69 FITC | H1.2F3 | BD Biosciences | cat# 553236 |
| Anti-mouse CD8a PerCP-Cy5.5 | 53-6.7 | BD Biosciences | cat# 551162 |
| Anti-mouse Ly6C BV421 | HK1.4 | Biolegend | cat# 128032 |
| Anti-mouse CD103 BV510 | M290 | BD Biosciences | cat# 563087 |
| Anti-mouse CD4 BV605 | RM4-5 | Biolegend | cat# 100548 |
| Anti-mouse CCR7 BV650 | 4B12 | BD Biosciences | cat# 564356 |
| Anti-mouse KLRG1 BV711 | 2F1 | BD Biosciences | cat# 564014 |
| Anti-mouse CD45<br>BUV395 | 30-F11 | BD Biosciences | cat# 564279 |
| Anti-mouse CD44<br>BUV737 | IM7 | BD Biosciences | cat# 612799 |
| Anti-mouse $\gamma\delta$ TCR PE | eBioGL3 (GL3) | Invitrogen | cat# 12-5711-82 |
| Anti-mouse CD3 PE-CF594 | 145-2C11 | Biolegend | cat# 100348 |
| Anti-mouse CD62L PE-Cy7 | MEL-14 | Invitrogen | cat# 25-0621-82 |
| Anti-mouse CD25 APC | PC61 | BD Bioscience | cat# 557192 |
| Anti-mouse CD11b AF700 | M1/70 | BD Biosciences | cat# 557960 |
| Anti-mouse FoxP3 AF488 | MF-14 | Biolegend | cat# 126406 |
| Anti-mouse RORγT<br>BV421 | Q31-378 | BD Bioscience | cat# 562894 |
| Anti-mouse $\gamma\delta$ TCR<br>BV510 | GL3 | BD Biosciences | cat# 563218 |
| Anti-mouse IL-17A<br>BV650 | TC11-18H10 | BD Biosciences | cat# 564170 |
| Anti-mouse NK1.1 BV711 | PK136 | BD Biosciences | cat# 740663 |
| Anti-mouse IFNγ BV785 | XMG1.2 | Biolegend | cat# 505837 |
| Anti-human/mouse T-bet<br>PE | eBio4B10 (4B10) | Invitrogen | cat# 12-5825-80 |
| Anti-mouse CD3 PE-CF594 | 145-2C11 | Biolegend | cat# 100348 |
| Anti-mouse Gata-3 PE-Cy7 | L50-823 | BD Biosciences | cat# 560405 |
| Anti-mouse IL-4 BV711 | 11B11 | BD Biosciences | cat# 564005 |

For intracellular staining, single cell suspensions were resuspended in 1 mL of Golgi Stop protein transport inhibitor diluted 1:1500 in RPMI with 10% FBS, 1% sodium pyruvate, 1X HEPES, 1X nonessential amino acid, 1% penicillin-streptomycin, phorbol 12-myristate 13-acetate (50 ng/ml), and ionomycin (1 µg/ml), and incubated for 4 hours at 37°C. Samples were washed once with FACS buffer and Fc receptors blocked with anti-mouse CD16/CD32. Samples were stained with antibodies listed in **Table 1** against surface markers and fixed and permeabilized with 1X fixation and permeabilization buffer and incubated at 4°C for 40 to 50 min protected from light. After incubation, samples were washed two times with 1X permeabilization and wash buffer from the transcription factor buffer kit (BD Biosciences) and stained with antibodies against IFN-γ, IL-17, IL-4, and the transcription factors RORγT, Gata-3, T-bet, and FoxP3 (**Table 1**), diluted in 1X permeabilization and wash buffer at 4°C for 40 to 50 min protected from light. Finally, samples were washed two times with 1X permeabilization and wash buffer and resuspended in FACS buffer. Total cell counts were determined by addition of counting beads to a known volume of sample after staining, just before cytometer acquisition.

For tetramer staining, biotinylated monomers H-2K(b) SIINFEKL and I-A(b) QAVHAAHAEIN were received from the NIH Tetramer Core Facility (Emory University, 954 Gatewood Road NE, Atlanta, GA 30329, USA) and tetramerized according to the NIH Tetramer Core Facility recommendations. Briefly, streptavidin was added every 10 minutes at room temperature 10 times and tetramers were conserved in the dark between each addition. The quantity of streptavidin added was calculated as described previously (*74*). PE-streptavidin and APC-streptavidin (cat# 4052454, cat# 405243, respectively BioLegend) were used to multimerize biotinylated monomers. Single cell suspensions from digested bladders were resuspended in 100 µl of FACS buffer containing tetramers diluted 1/100 and incubated 1 hour at 4°C in the dark. We used a double staining approach, in which the same tetramer was labeled with APC or PE and both preparations were used in the same samples. After tetramer staining, cells were incubated with blocking anti-mouse CD16/CD32 and antibodies against cell surface markers as described above

### Immune cell depletion

To deplete T cells (**Fig. 1**), 100 µg of CD4 or 100 µg of CD8 (clone GK1.5, clone YTS 169.4, respectively, Bio X Cell) depleting antibodies were injected intraperitoneally in 100 µL sterile PBS per mouse 3 days before primary infection and repeated 7 days after the first injection, *i.e.*, day 4 PI. To deplete CD4^+^ and CD8^+^ T cells (**Fig. 6**) 100 µg of CD4 and 100 µg of CD8 depleting antibodies were injected intraperitoneally in 100 µL sterile PBS per mouse 24 hours before challenge infection. 100 µg (**Fig. 1**) or 200 µg of the isotype control (**Fig. 6**) (clone LTF-2, Bio X Cell) were injected intraperitoneally in 100 µL of sterile PBS per mouse at the same time in control groups.

### Intravascular labeling

Female mice were anesthetized as describe above, and 3µg of Pacific Blue anti-CD3 (clone 500A2, BD bioscience), or 3µg of PE anti-CD3 (clone 145-2C11, BD bioscience) were injected intravenously in the retroorbital sinus. 3 minutes later mice are sacrificed as described above, blood and bladder are collected and analyzed by flow cytometry or by confocal microscopy.

### Histological and immunostaining for confocal microscopy

Whole bladders were fixed with 4% paraformaldehyde (PFA) in PBS for 1 hour and subsequently washed with PBS. Samples were then dehydrated in 30% sucrose in PBS for 24 hours. Samples were cut transversally and embedded in optimal cutting temperature compound, frozen, and sectioned at 30 µm. Sections were blocked for 1 hour with blocking buffer [3% bovine serum albumin (BSA) + 0.1% Triton X-100 + goat serum (1:20) in PBS] and washed three times. Immunostaining was performed using rat anti-mouse CD3 (1:100) in staining buffer (0.5% BSA + 0.1% Triton X-100 in PBS) overnight. Sections were washed and stained with phalloidin (1:350), goat anti-hamster (1:350) and goat anti-rat (1:1000) antibodies in staining buffer for 4 hours. Last, sections were washed and stained with 4′,6- diamidino-2-phenylindole. Confocal images were acquired on a Leica SP8 confocal microscope. Final image processing was done using Fiji (version 2.0.0-rc-69/1.52p).

### Adoptive T cell transfer

To purify T cells from infected and resolved mice, spleens and bladder draining lymph nodes were gently pressed through a 70µm filter, pooled, and counted. CD3^+^ cells were isolated with the Pan T Cell Isolation Kit II (Miltenyi) using magnetic separation (negative selection) according to manufacturer’s recommendations. CD3^+^ T cells were counted and diluted in PBS to 10^7^ cells/ml or 10^6^ cells/ml and 100µl were injected intravenously in naïve mice via the retroorbital sinus.

### FTY720 treatment

Female mice received by an intraperitoneal injection 50µl of aqueous solution, or FTY720 at 0,4mg/ml diluted in aqueous solution (0,02mg/mouse ≈ 1mg/kg) at 3 consecutive days (day −2, −1, and at the time of challenge infection).

### Statistical analysis

Statistical analysis was performed in GraphPad Prism 9 (GraphPad, USA) for Mac OS X applying the nonparametric Mann-Whitney test for unpaired data in the case of two group comparisons. To correct for comparisons made within an entire analysis or experiment, calculated *p*-values were corrected for multiple testing with the false discovery rate (FDR) method to determine the FDR-adjusted *p*-value. In the case that more than two groups were being compared, Kruskal-Wallis tests were performed, with Dunn’s post hoc test to correct for multiple comparisons. All adjusted *p*-values are shown in the figures.

## Acknowledgments

We are thankful for insightful discussion with Dr. Elizabeth Wohlfort, David Withers, and Bruno Lucas. We thank the NIH Tetramer Core Facility (contract number 75N93020D00005) for providing H-2K(b) SIINFEKL, and I-A(b) QAVHAAHAEIN biotinylated monomers used in this study. Funding: LLM was part of the Pasteur-Paris University (PPU) International PhD Program, which received funding from the European Union’s Horizon 2020 research and innovation program under the Marie Sklodowska-Curie grant agreement no. 665807 and from the Labex Milieu Intérieur (ANR-10-LABX-69-01). MAI was supported by funding from the *Agence Nationale de la Recherché* (French National Research Agency) ANR-17-CE17-0014 and ANR-19-CE15-0015 and *Chaires d’excellence de l’IdEx Université de Paris*.

## Author contributions

Conceptualization: MR and MAI; Methodology: MR, LLM, TC, MAI; Investigation and data analysis: MR, LLM, TC, MAI; Writing - Original Draft: MR and MAI; Writing - Review & Editing: MR, LLM, TC, MAI; Funding Acquisition: MAI; Supervision: MAI.

## Competing interests

The authors declare no competing interests.

## Data and materials availability

Further information and requests for resources and reagents should be directed to and will be fulfilled by Matthieu Rousseau and Molly A. Ingersoll.

## List of Supplementary data

**Supplementary Figure 1:** T cell infiltration

**Supplementary Figure 2:** No distinct CD4^+^ Th cell bias arises during primary and recurrent UTI in the lymph nodes

**Supplementary Figure 3:** Limiting antigen persistence does not change Th cell polarization in bladder-draining lymph nodes

**Supplementary Figure 4:** Limiting antigen persistence does not impact bladder-draining lymph node T cells with a tissue resident memory phenotype

## Supplementary Information for

**Supplementary Figure 1:**
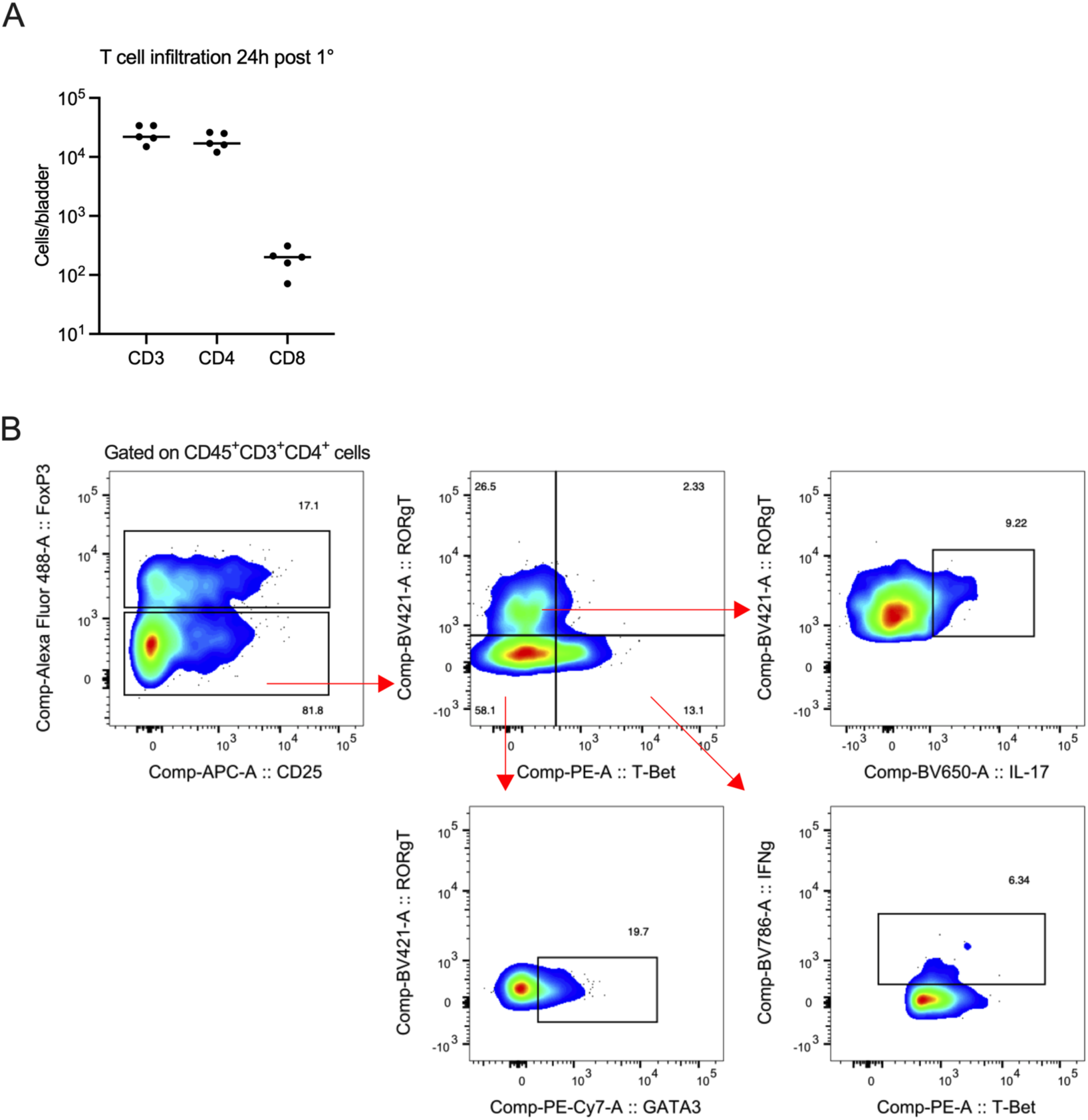
T cell infiltration. (**A**) Graph shows the total number of the specified T cell populations per bladder 24 hours post primary infection. A representative experiment from 3 experiments is presented. Each circle is a mouse and lines are medians. (**B**) Representative flow cytometry plots show the gating strategy to determine T cell infiltration and polarization by transcription factor and cytokine expression after primary of challenge infection. Bladder is shown and the same strategy was used for lymph nodes shown in **Fig. S2** and **S3**.

**Supplementary Figure 2:**
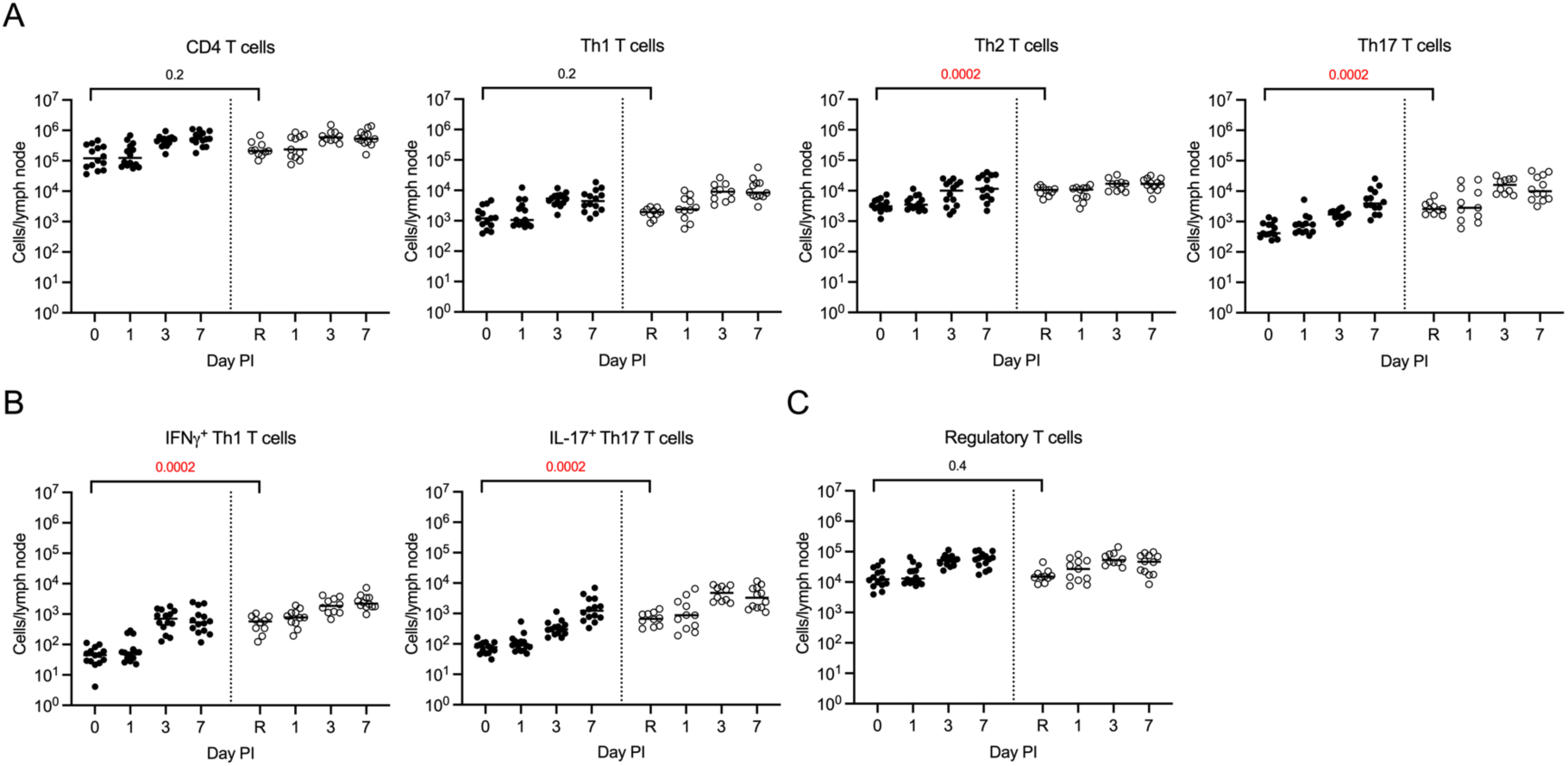
No distinct CD4^+^ Th cell bias arises during primary and recurrent UTI in the lymph nodes. Six-week-old infected female C57BL/6 mice were sacrificed at the indicated timepoints post-primary (filled circles), or post-challenge infection (open circles). Day 0 are naïve mice and ‘R’ denotes animals that resolved their primary UTI but were not challenged with a second infection. The graphs depict total (**A**) CD4^+^, Th1, Th2, or Th17 T cells, (**B**) IFNg^+^ Th1 or IL-17^+^ Th17 T cells, and (**C**) Tregs, in the draining lymph nodes at the indicated day post primary or challenge infection. Gating strategy is shown in **Fig. S1B**. Data are pooled from 2 experiments, n=5 to 7 mice/group in each experiment. Results from the bladders of these mice are shown in **Fig. 2**. Each circle is a mouse and lines are medians. Nonparametric Mann-Whitney tests comparing T cell accumulation between ‘0’ and ‘R’ were performed. All *p*-values were corrected for multiple comparisons across all populations shown using the false discovery rate (FDR) method and *p*-values <0.05 are in red.

**Supplementary Figure 3:**
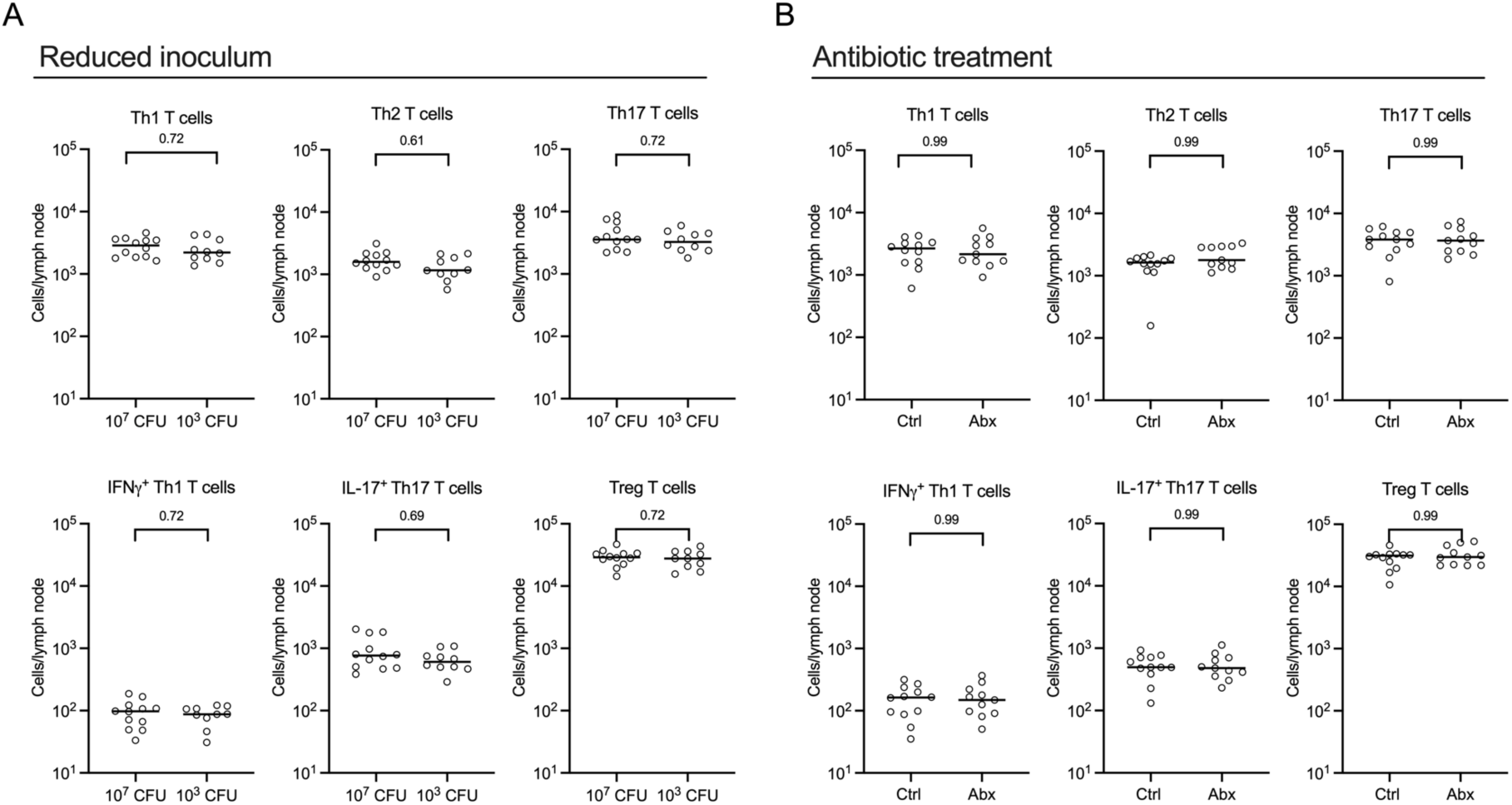
Limiting antigen persistence does not change Th cell polarization in bladder-draining lymph nodes. Graphs show the total number of specified cell populations per bladder-draining lymph nodes from mice presented in **Fig. 4A-D** in which (**A**) shows draining lymph nodes of mice presented in **Fig. 4A** and **B, (B**) shows draining lymph nodes of mice presented in **Fig. 4C** and **D,** analyzed by flow cytometry. Gating strategy is shown in **Fig. S1B**. Data are pooled from 2 experiments, n=4 to 6 mice/group in each experiment (**A**), or n=5 to 6 mice/group in each experiment (**B**). Each circle is a mouse and lines are medians. Nonparametric Mann-Whitney tests comparing cell numbers between each condition were performed. *p*-values were corrected for multiple comparisons using the false discovery rate (FDR) method. Corrected *p*-values are shown.

**Supplementary Figure 4:**
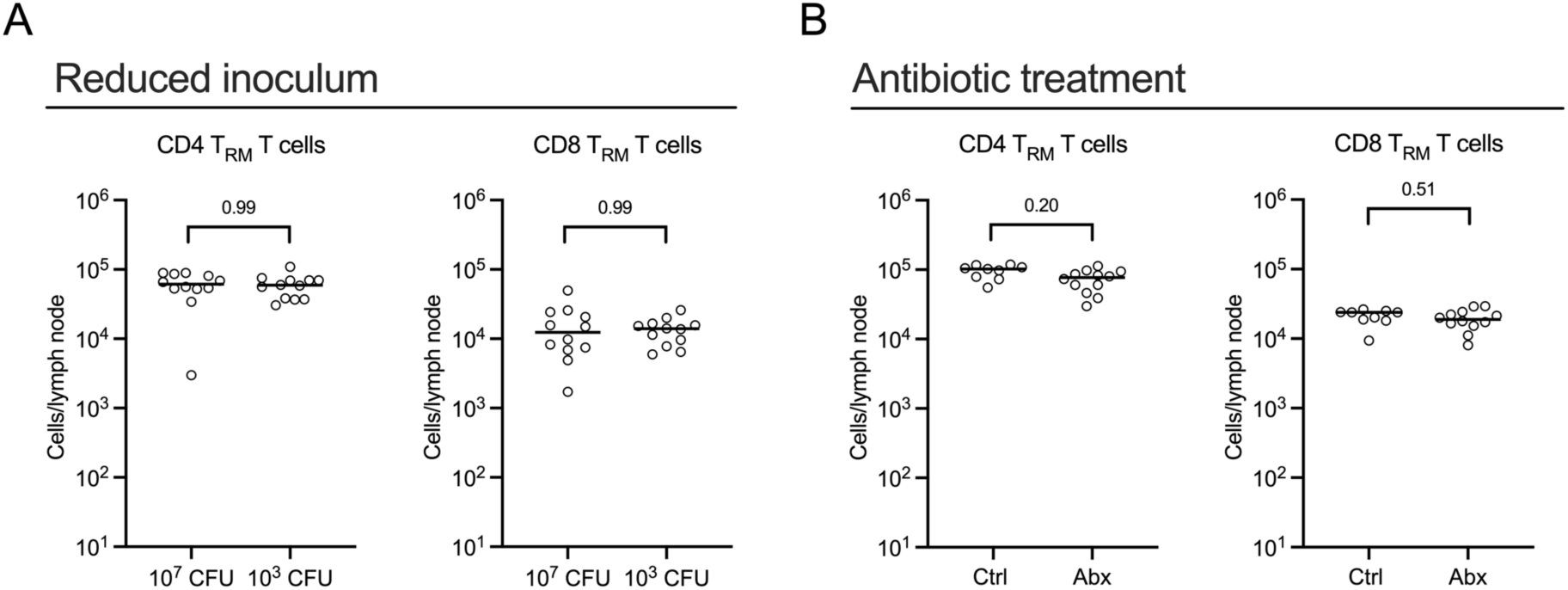
Limiting antigen persistence does not impact bladder-draining lymph node T cells with a tissue resident memory phenotype. (**A, B**) Graphs show the total number of specified cell populations per bladder draining lymph nodes from mice (**A**) presented in **Fig. 5A** and (**B**) presented in **Fig. 5B**. Data are pooled from 2 experiments, n=3 to 6 mice/group in each experiment. Nonparametric Mann-Whitney tests comparing cell numbers between each condition were performed. *p*-values were corrected for multiple comparisons using the false discovery rate (FDR) method. Corrected *p*-values for each comparison are presented.

## Notes

### Competing Interest Statement

The authors have declared no competing interest.

### Summary of Updates

Additional data included

